# Pasta, a versatile transcriptomic clock, maps the chemical and genetic determinants of aging and rejuvenation

**DOI:** 10.1101/2025.06.04.657785

**Authors:** Jérôme Salignon, Maria Tsiokou, Patricia Marqués, Enriqueta Rodríguez-Diaz, Hazel Ang, Federico Pietrocola, Christian G. Riedel

## Abstract

With the growing burden of age-related diseases, understanding and modulating the aging process has become a priority. Transcriptomic aging clocks (TACs) can track biological age but remain limited by platform dependence, tissue specificity, or restricted accessibility. To address this, we developed Pasta, a robust and broadly applicable human TAC built using a novel ‘age-shift’ learning framework. Pasta accurately predicted relative age across diverse tissues, data types, including bulk and single-cell RNA-Seq as well as microarray data, and species. Its predictions aligned with senescent and stem-like cellular states, with model coefficients enriched for p53 and DNA damage response pathways. Pasta’s age scores correlated with tumor grade and patient survival in several cancer types, indicating potential clinical relevance. Applied to the Connectivity Map L1000 dataset, Pasta identified both established and previously unrecognized age-modulatory compounds and genetic perturbations, highlighting mitochondrial translation and mRNA splicing as key determinants of cellular propensity for aging and rejuvenation, respectively. Experimental validation confirmed pralatrexate as a potent senescence inducer and piperlongumine as a rejuvenating agent. Together, these findings establish Pasta as a versatile and accessible tool for aging research and therapeutic discovery.

## Introduction

The growing prevalence of chronic non-communicable diseases in later life has positioned aging as a major driver of pathology. This, coupled with advances in understanding the fundamentals of aging and health^1,2^, has intensified interest in pharmacological and lifestyle strategies to modulate aging^3^. Unlike chronological aging, which progresses at a constant pace, biological aging is modulated by endogenous and environmental factors that can accelerate, decelerate, or even reverse aging processes^4^. This highlights biological age as a latent but measurable trait that reflects current and future health, captures individual variability in aging rates, and offers a means to evaluate geroprotective interventions.

Over the past decade, a range of computational tools—collectively termed ‘aging clocks’—have been developed to quantify biological age and estimate aging rates. These models draw from diverse molecular and clinical hallmarks across multiple ‘omics’ layers, including epigenetic, transcriptomic, metabolomic, proteomic, and metagenomic data^5^. Among them, epigenetic clocks based on genome-wide CpG methylation patterns have been widely used due to their high precision in estimating chronological and biological age, strong predictive value for morbidity and mortality, and the relative stability of such DNA methylation (DNAm) marks, which enable the capture of cumulative aging effects^6^. However, DNAm clocks also have limitations, including reduced sensitivity to rapid or transient changes^7^ and limited biological interpretability, as CpGs are not always clearly linked to gene function^8^. In addition, single-cell DNAm approaches remain costly, technically demanding, and lack standardized protocols, restricting their utility in resolving aging at single-cell resolution^6^.

Transcriptomic aging clocks (TACs) address these limitations: they sense transient shifts, yield interpretable gene and pathway outputs, and can be applied to extensive bulk^9^, single-cell^10,11^, spatial^10,12^, genetic^13,14^ and chemical^13,15^ perturbation datasets. Notably, ongoing efforts include constantly growing transcriptomic perturbation resources, such as the Connectivity Map (CMAP)L1000^13^, Perturb-Seq datasets^14^ or the Tahoe-100 dataset^15^. Applying TACs to such datasets could reveal regulators of aging and rejuvenation with translational potential. For example, compounds that raise the biological age of cancer cells may enhance therapeutic efficacy^16^, and gene perturbations that accelerate aging in iPSC-derived neurons can improve models of late-onset disease^17^. Conversely, agents that lower age in normal cells may promote regeneration; improving stem cell fitness^18^, iPSC protocols^19^, or partial reprogramming cocktails^20^. While a handful of age-modulatory perturbations have recently been described^21^, systematic approaches to identify them remain limited. A recent study showed the potential of this approach by mining the Gene Expression Omnibus (GEO)^22^ database using aging clocks to find novel age-modulatory perturbations^23^. A landmark and untaped resource in this context is the CMAPL1000 dataset^13^, which contains over three million transcriptomes from 248 cell lines exposed to more than 14,000 gene perturbations and 30,000 compounds. Identifying both general and cell line-specific age-modulatory perturbations in this dataset could inform the development of more effective age-related translational strategies.

Human TACs have been developed in blood^24^, muscle^25^, fibroblasts^26^, skin^27^, and multi-tissue settings^28–32^, and more recently at single-cell resolution using blood^33^, immune cells^34,35^, and PBMCs^36^. Despite this progress, no TAC has achieved widespread adoption comparable to epigenetic clocks, due to limitations in performance, availability, and platform compatibility. Gene expression sensitivity to environmental cues enables detection of transient states but also introduces variability and noise, requiring robust modeling across heterogeneous datasets. Most TACs, including single-tissue models and the multi-tissue RNAAgeCalc^29^, are trained on single datasets, leading to overfitting and limited generalization. Consistently, MultiTIMER^30^, trained on multiple datasets, outperforms RNAAgeCalc^29^ despite being trained on fewer samples (∼3,000 vs ∼9,600). Another key barrier is availability: DeepQA^32^ and Shokhirev & Johnson^28^’s TACs offer no usable software, while RAPToR^31^ requires users to generate custom references without default human datasets. Moreover, most multi-tissue TACs^28–30,32^ require raw data reprocessing, batch correction, or model retraining, limiting their use to bulk RNA-Seq data and reducing reproducibility.

To address these limitations, we present Pasta (Predicting age-shift from transcriptomic analysis), a multi-platform, multi-tissue, ready-to-use TAC. It outperformed MultiTIMER^30^, the currently best available multi-tissue human TAC, across diverse datasets, yielded meaningful predictions across tissues and platforms in mouse datasets, and had its predictions being largely influenced by p53-related genes. Beyond age prediction, Pasta effectively distinguished between senescent, quiescent, and stem cells, in both static and dynamic settings, indicating its capacity to capture bidirectional cell state transitions from stemness to senescence. This capability could be valuable in clinical contexts where Pasta may help to predict tumor grade. When applied to the CMAP L1000 dataset^13^, Pasta highlighted potential age modulators such as pralatrexate and piperlongumine, which were experimentally validated, and pinpointed molecular features associated with cell-line responsiveness to pro-or anti-aging interventions. Taken together, Pasta offers a robust and interpretable framework for biological age estimation across tissues, platforms, and species, and for systematic discovery of genetic or chemical modulators of aging and rejuvenation, with potential relevance to translational research in oncology^16^, neurodegeneration^17^, and regenerative medicine^18–20^.

## Results

### Construction of a general-purpose transcriptomic aging clock from heterogeneous multi-tissue data

Given that existing TACs often show limited transferability, being trained on single cohorts or requiring extensive preprocessing, we aimed to develop a broadly applicable transcriptomic aging clock that can be used across tissues and sequencing platforms. To ensure broad generalization, we compiled heterogeneous transcriptomic data from the Genotype-Tissue Expression (GTEx)^37^ project, the GEO, and the Expression Atlas^38^, retaining 17,212 healthy samples spanning multiple tissues from 21 studies (18 bulk RNA-Seq and 3 microarray; Fig. 1a, Supplementary Table 1). Gene expression values were rank-transformed to mitigate platform and processing effects.

**Figure 1.**
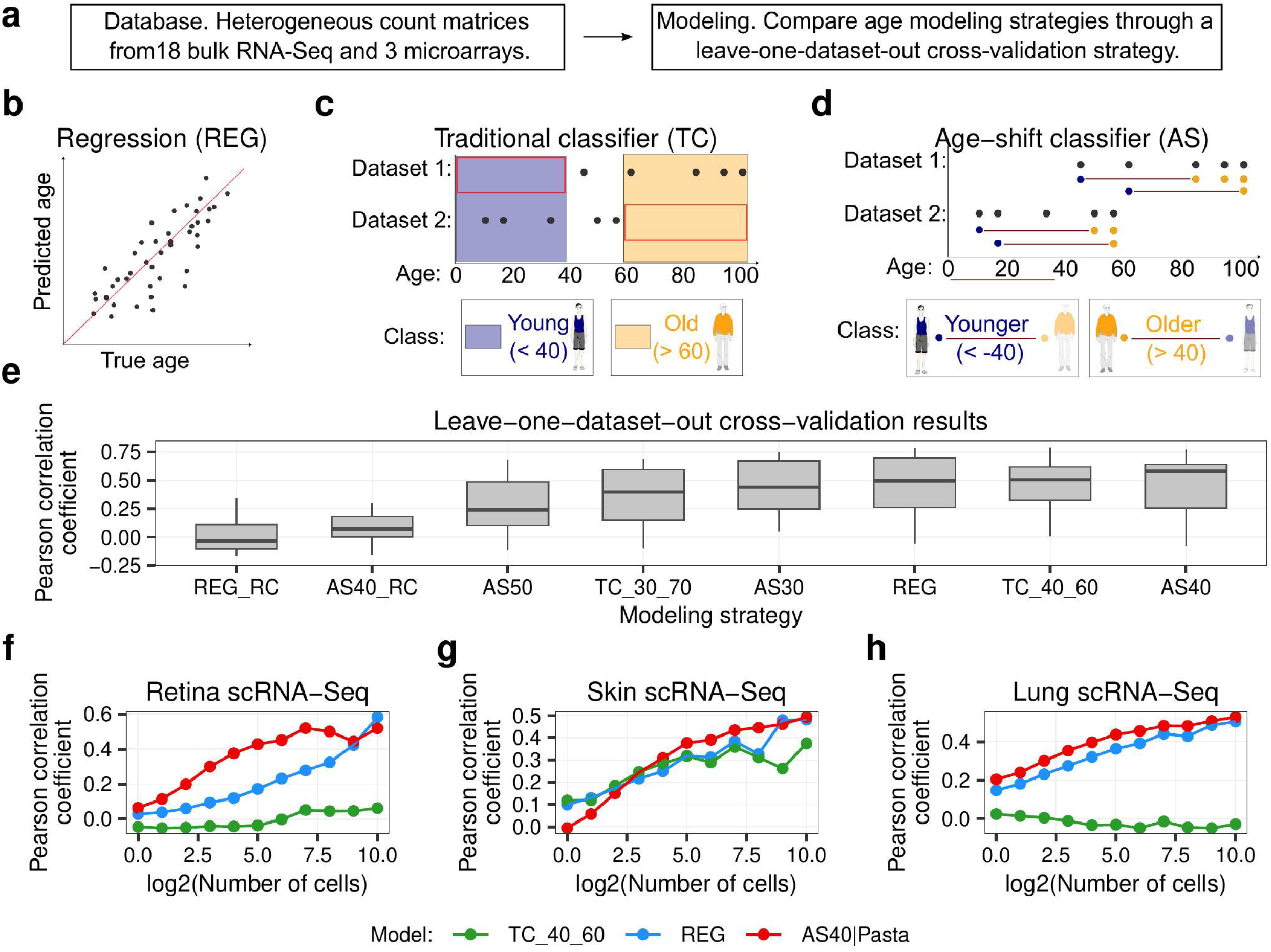
The age-shift classifier enables accurate cross-platform prediction of relative transcriptomic age. (**a**) Overview of the evaluated modelling strategies. Models were trained and tested exclusively on samples from healthy donors. (**b-d**) Schematic diagrams of the regression (**b**), traditional classifier (**c**), and age-shift classifier (**d**) strategies. (**e**) Comparative performance in predicting relative age, assessed using leave-one-dataset-out cross-validation. All samples were rank transformed except for REG_RC and AS40_RC where raw counts were used. The y-axis shows the Pearson Correlation Coefficient (PCC) between chronological ages and predicted age scores in the held-out datasets. (**f-h**) Age predictions for single-cell RNA-Seq aging atlases of the retina^44^ (**f**), skin^45^ (**g**), and lung^43^ (**h**). Abbreviations: REG, regression; AS30/40/50, age-shift classifier with 30, 40, or 50 year age difference; TC_30_70 and TC_40_60, traditional classifiers with young cutoffs at 30 or 40 years and old cutoffs at 70 or 60 years; RC, raw counts.

Using this dataset, we compared three modeling strategies: regression models predicting chronological age (Fig. 1b), young-versus-old classifiers (Fig. 1c), and age-shift classifiers trained on sample pairs separated by a defined age gap (Fig. 1d) – the latter being an approach conceptually similar to twin neural network regression^39,40^ and pairwise difference regression^41^ or classification^42^. For the classifiers, predictions were converted to age scores (see Methods).

We evaluated all models using a leave-one-dataset-out (LODO) strategy. An age shift model with a 40-year age gap achieved the highest median Pearson correlation coefficient (PCC) across all samples (Fig. 1e) and remained the top performer when restricting the evaluation to young and old individuals only (Extended Data Fig. 1a). It also showed consistent performance across tissues and platforms (Extended Data Fig. 1b, 2, and 3). Models trained on unranked counts performed poorly (median PCC < 0.1), underscoring the importance of rank transformation (Fig. 1e; Extended Data Fig. 1a).

To assess generalization to data types unseen during training, we tested our top models on single-cell RNA-Seq datasets. Because training relied on bulk data, single-cell profiles were aggregated into pseudobulks of increasing size (1 to 1024 cells) to evaluate performance at different resolutions. Across retina, skin, and lung datasets, the age-shift model consistently outperformed the regression model and the young-versus-old classifier (Fig. 1f–h)^43–45^. Notably, the latter failed when test datasets lacked age categories present during training (Fig. 1f,h), highlighting the limitation of fixed age cutoffs.

Altogether, the 40-year age-shift model provided the most robust and generalizable predictions of relative age across heterogeneous transcriptomic data, outperforming previous approaches. We refer to this model as Pasta throughout the remainder of this study.

### Pasta leverages p53-related genes to accurately predict relative age across platforms, tissues, and species

To evaluate Pasta’s performance, we compared it with MultiTIMER, the most accurate available multi-tissue human TAC^30^. Pasta outperformed MultiTIMER in 7 of 8 datasets from our LODO analysis (Fig. 2a; Supplementary Table 2) and in 5 of 6 independent validation datasets (Fig. 2b). These datasets encompassed diverse tissues (Supplementary Table 2), confirming Pasta’s broad applicability across tissue types. Since Pasta works across tissues and platforms, we wondered if it could also work across species. We extended Pasta for usage in mouse by leveraging one-to-one orthologues (see Methods). Encouragingly, we observed that Pasta could also predict age accurately in mouse across a variety of tissues and platforms (Fig. 2c,d). Altogether these results show that Pasta is an accurate predictor of relative age across tissues, platforms, and species.

**Figure 2.**
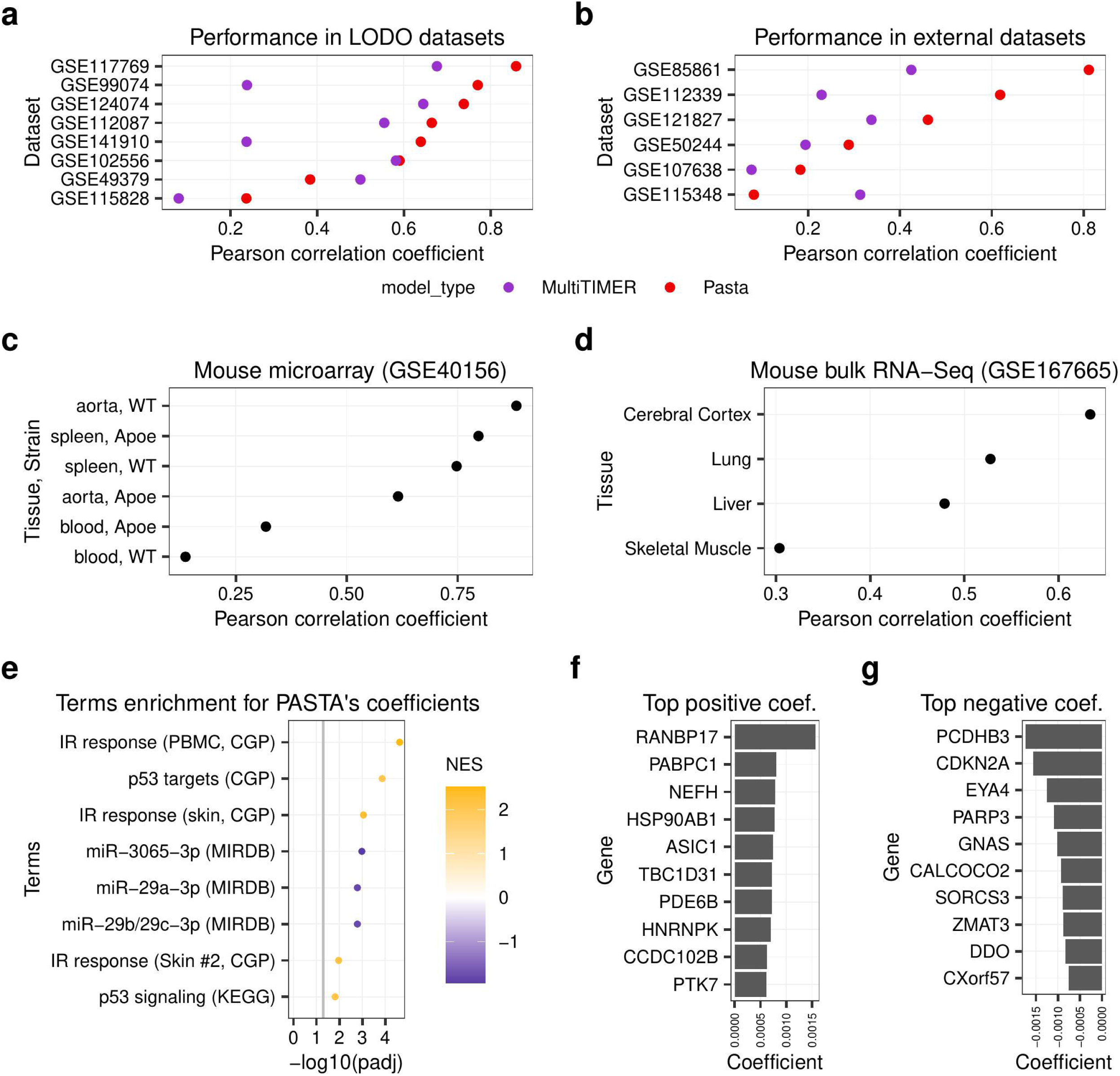
Pasta predicts transcriptomic age across tissues through p53-linked signals. (**a-b**) Pearson Correlation Coefficient (PCC) of age scores for Pasta and MultiTIMER compared with chronological ages for healthy individuals. (**a**) Results from the leave-one-dataset-out analysis. (**b**) Results on external validation datasets. (**c-d**) PCC of age scores for Pasta compared with chronological age for mouse datasets GSE40156 (**c**) and GSE167665 (**d**). (**e**) Gene set enrichment analysis (GSEA) of Pasta coefficients using the REACTOME, CGP, KEGG and MIRDB gene sets from the Human MSigDB Collections. The grey lines mark the adjusted p-value cutoff of 0.05. (**f-g**) Top ten genes with the most positive (**f**) and negative (**g**) coefficients.

We next conducted <u>g</u>ene <u>s</u>et <u>e</u>nrichment <u>a</u>nalysis (GSEA) on broadly used collections (REACTOME, miRDB…)^46^ using Pasta’s coefficients. Genes with positive coefficients, whose higher expression predicted older age, were enriched for p53 signaling and irradiation response terms, pointing to activation of the DNA damage-p53 axis^47^ (Fig. 2e; Supplementary Table 3). The second and third strongest negative enrichments, genes whose higher expression predicted younger age, were targets of miR29, a p53 activator^48^ which drives aging phenotypes in mice^49^. Notably, no pathway was significantly enriched when using the baseline regression model generated in Figure 1, suggesting that the age-shift learning framework more effectively captures aging-relevant transcriptional signal.

Consistent patterns were observed at the individual gene level, with RANBP17 carrying the largest negative coefficient, nearly twice that of any other gene (Fig. 2f). This gene promotes cell proliferation^50^ and has been proposed to be a master regulator of aging which decreases with age^51^. The strongest positive coefficients belonged to PCDHB3, a colorectal-cancer tumour-suppressor^52^, CDKN2A (p16), a canonical p53 target and senescence marker, and ZMAT3, another key p53 target^53^ (Fig. 2g).

In conclusion, Pasta is a robust multi-tissue, multi-platform, and multi-species transcriptomic age model which grounds its predictions in tumor-suppressor genes and core aging modulators.

### Pasta accurately tracks cellular age across senescence, pluripotency and differentiation states

Given Pasta’s strong performance across tissues, we then asked whether it could also track cellular aging in vitro, focusing on senescent and stem cells, which represent opposite ends of the cellular age spectrum.

Pasta accurately distinguished senescent from proliferative cells in 19 of 30 RNA-Seq datasets (AUC-ROC = 1) and achieved moderate discrimination (AUC-ROC > 0.7) in five additional datasets (Fig. 3a; Supplementary Table 4), consistently labeling senescent cells as older. It also distinguished senescent from quiescent cells in 4 of 5 datasets (Fig. 3b). In a 450-day serial passaging experiment of primary fibroblasts that reached replicative senescence, Pasta’s age scores increased almost linearly with culture duration (PCC = 0.896; Fig. 3c). We next assessed whether Pasta could detect chemical inducers of senescence in single-cell data. Combining two studies that tested the same 138 compounds in A549 cells, one using scRNA-Seq^54^ and another using flow cytometry with a senescence reporter^55^, Pasta assigned significantly higher age scores to senescence-inducing compounds than to other compounds (*p* = 9.9 × 10^−6^; Welch’s two-sample t-test; Fig. 3d).

**Figure 3.**
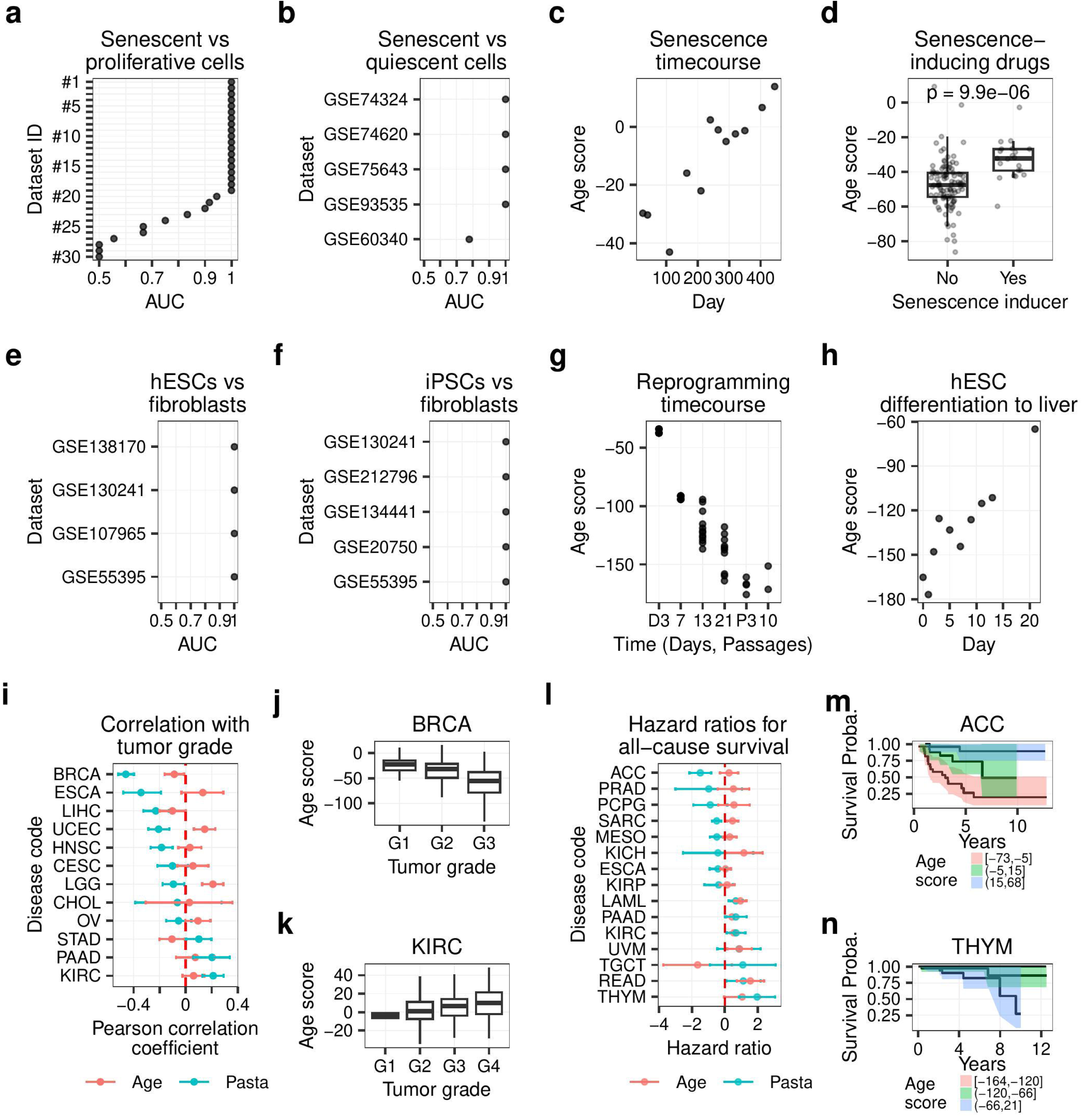
Pasta identifies senescent and stem cells and may help stratify tumor grade and patient overall survival. (**a-b**) Pasta’s ability to discriminate between senescent and proliferative (**a**) or quiescent (**b**) cells, assessed using AUC-ROC (Area Under the Curve of the Receiver Operating Characteristic) values, across multiple RNA-Seq datasets. (**c**) Pasta’s predictions on a microarray senescence time course dataset^129^. (**d**) Pasta was applied to pseudobulks from a SciPlex dataset generated by exposing the A549 cell line to 138 drugs^54^, and the resulting age scores were evaluated for their capacity to recover senescence-inducing drugs, as reported in Wang et al 2017^55^. Only the highest drug concentration (10 µM) was used for this analysis. (**e-f**) Pasta’s ability to discriminate between stem cells (**e**) or induced-pluripotent stem cells (iPSCs) (**f**) and fibroblasts, assessed by AUC-ROC. (**g**) Pasta’s age scores on a bulk RNA-Seq OSKM-mediated reprogramming time-course dataset^130^. (**h**) Pasta’s age scores on a bulk RNA-Seq liver differentiation time-course dataset^131^. (**i**) Correlations of patient chronological ages (red) and Pasta’s age scores (blue) with tumor grade. (**j-k**) Boxplots of Pasta’s age scores for breast invasive carcinoma (**j**) and kidney renal clear cell carcinoma (**k**) patients, grouped by tumor grade. (**l**) Hazard ratios from Cox proportional-hazards regression models fitted for scaled ages (red) and scaled age scores (blue). (**m**-**n**) Kaplan-Meier survival estimates for adrenocortical carcinoma (**m**) and thymoma (**n**) patients, grouped by age score tertiles.

Having established that Pasta can detect senescent and thus very old cells, we next tested its sensitivity to very young cell states. Pasta effectively distinguished embryonic stem cells from fibroblasts across four bulk RNA-Seq datasets (Fig. 3e) and induced pluripotent stem cells from fibroblasts in five datasets (Fig. 3f), consistently labeling stem cells as younger. During OSKM-mediated fibroblast reprogramming, Pasta’s age scores declined linearly with time (PCC = −0.905; Fig. 3g), whereas in directed differentiation of human embryonic stem cells toward liver cells, age scores increased steadily (PCC = 0.926; Fig. 3h).

Together, these results show that Pasta reliably tracks not only tissue age but also cellular age across a continuum of states, from senescence to pluripotency and differentiation.

### Pasta reveals cancer type specific associations between cellular age, tumor grade, and patient survival

Studies using transcriptomic stemness models have shown that higher stemness correlates with advanced tumor grade and poorer survival in cancer patients^56,57^. Recently, increased cancer cell age has also been linked to adverse outcomes^16^. Indeed, the accumulation of senescent cells within tumors, whether normal or cancerous, can promote inflammation, relapse, and immune evasion^16,58–60^. Yet, data from The Cancer Genome Atlas (TCGA), the largest transcriptomic cancer resource, have not been systematically examined using an aging clock, leaving it unclear whether poor prognosis is driven by high stemness alone or also by senescence.

To test this, we applied Pasta to TCGA tumors, leveraging its capacity to identify both youthful (high-stemness) and aged (senescent-like) transcriptional states. Pasta’s score showed weak correlation with donor age (PCC=0.0570), so unadjusted values were used in subsequent analyses. Six of twelve grade annotated cancer types showed significantly negative correlations between age scores and tumor grade (Fig. 3i, j, Supplementary Table 5), while two**—**kidney renal clear cell carcinoma (KIRC) and pancreatic adenocarcinoma (PAAD)**—**showed significantly positive correlations (Fig. 3i, k). These results indicate that both increased and decreased biological age can be associated with higher tumor grade, consistent with roles of both stemness and senescence in tumor progression. Donor age significantly predicted grade in only two cancers (Lower Grade Glioma, LGG, and Uterine Corpus Endometrial Carcinoma, UCEC). Pasta’s stronger associations with tumor grade than donor age suggest that its scores capture biological rather than chronological age.

To test whether Pasta’s age scores were also linked to clinical outcomes, we fitted Cox proportional hazards models across 33 cancer types. After multiple testing correction, younger age scores were significantly associated with poorer survival in Adrenocortical carcinoma (ACC), Sarcoma (SARC), and Skin Cutaneous Melanoma (SKCM) (Fig. 3l,m; Supplementary Table 6), whereas older age scores were significantly associated with poorer survival in Thymoma (THYM) and Acute Myeloid Leukemia (LAML) (Fig. 3l,n). Several other cancers showed strong associations. Chronological age was a weaker predictor of survival in 18 cancer types and was uniformly associated with poorer outcomes.

Taken together, Pasta outperformed chronological age as a predictor of tumor grade and survival in several cancer types.

### Systematic identification of age-increasing and rejuvenating small molecules in the CMAP L1000 dataset

Having established Pasta’s predictive power for tissue and cellular aging in health and disease, we next assessed whether it can also identify age-modulatory perturbations.

To this end, we applied Pasta to the CMAP L1000 dataset^13^, which contained the transcriptomic responses of 248 cell lines to more than 30,000 chemical and 14,000 genetic perturbations, totaling more than three million transcriptomes. Coincidentally, the bead-based nature of the L1000 assay provided also an opportunity to test Pasta’s cross-platform robustness. Age scores were calculated for each perturbation and plate-normalized to corresponding negative controls (DMSO for compounds, empty vectors for genetic perturbations).

By this approach, we identified 259 compounds that were significantly associated with higher age scores and 59 compounds with lower age scores, hereafter termed *Aging* and *Rejuvenating* compounds, respectively (Fig. 4a; Supplementary Table 7). As expected, top Aging compounds included canonical senescence inducers such as mitoxantrone, gemcitabine, and doxorubicin^61^, whereas top rejuvenating compounds included pluripotency-promoting molecules, such as RepSox and PD0325901^62^. Annotating the CMAP library with known senescence-^61^ and reprogramming-^62^ related drug classes revealed that 18.9% of Aging compounds belonged to senescence-inducing classes, compared with less than 0.2% among all remaining compounds (*p* = 5.45 × 10^−76^, Fisher’s exact test, Fig. 4b). Similarly, 16.9% of Rejuvenating compounds belonged to reprogramming-related classes, compared with less than 0.2% among all remaining compounds (*p* = 3.01 × 10^−18^, Fisher’s exact test). Notably, 27.1% of Rejuvenating compounds were annotated as both pro-senescence and pro-reprogramming, all of which were histone deacetylases (HDAC) inhibitors, a drug class known to induce both reprogramming^63,64^ and senescence^65–67^.

**Figure 4.**
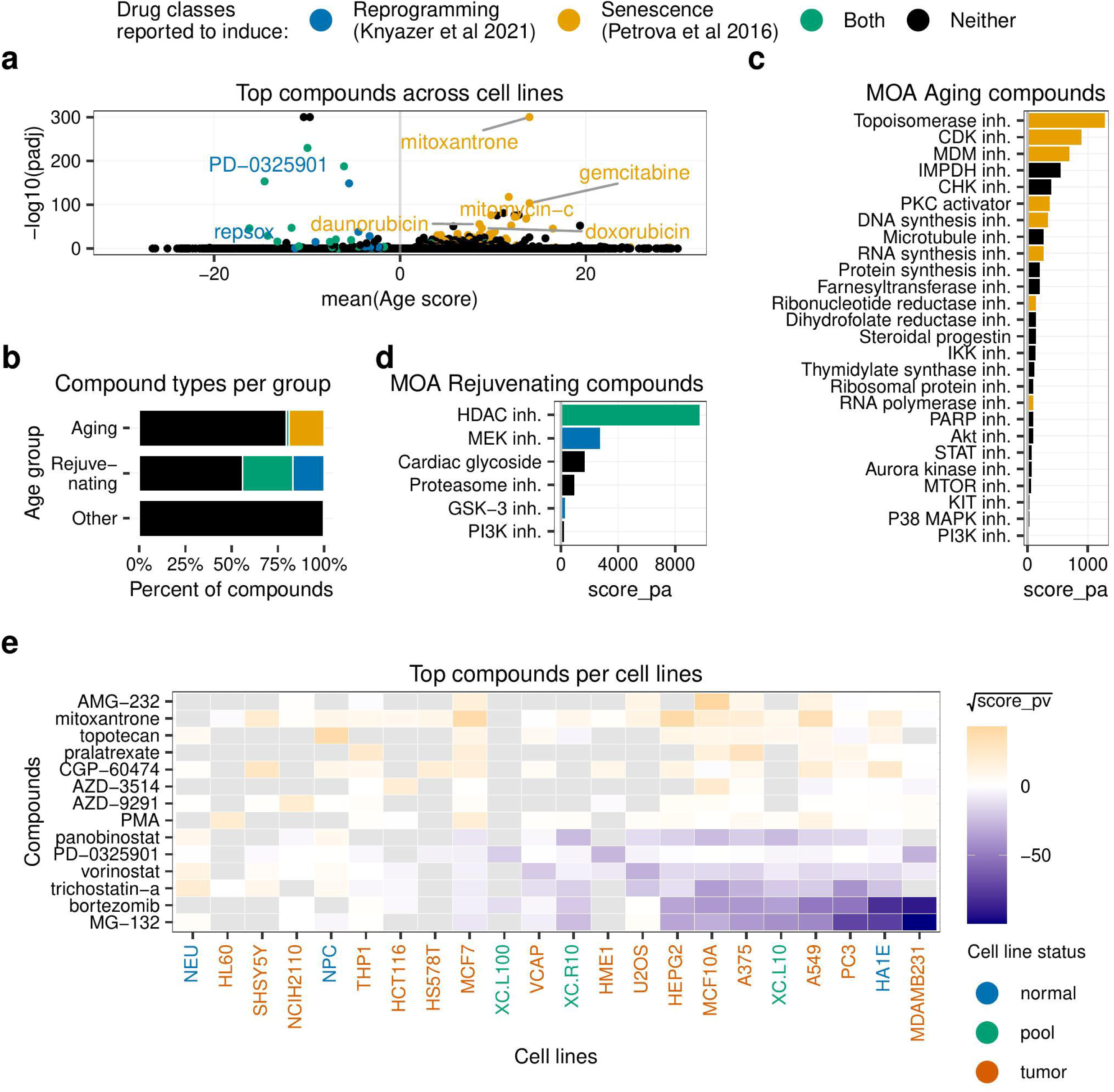
Pasta identifies age-modulatory compounds. (**a-d**) Yellow, blue, green, and black indicate drug classes or compounds from drug classes that induce senescence^61^, promote reprogramming^62^, affect both processes, or neither, respectively. (**a**) Volcano plot showing the most significant Rejuvenating compounds with negative age scores and Aging compounds with positive age scores. For display purposes, p-values equal to zero were set to 10^−300^. (**b**) Proportion of compounds annotated as senescence-inducing, pro-reprogramming, both, or neither for significantly age-increasing (Aging), age-decreasing (Rejuvenating), and other (Other) compounds. (**c-d**) Significantly enriched mechanisms of action (MOA) for Aging (**c**) and Rejuvenating (**d**) compounds. The grey lines mark the adjusted p-value cutoff of 0.05. (**e**) Heatmap of compounds by cell line showing the most significant age scores (see Methods). Colors indicate the square root of a metric balancing significance and effect size: *-log_10_(adjusted p-value) × mean(age score)*. Missing values are shown in grey.

When repeating the analysis using the baseline regression model from Figure 1, we found the regression model to be inferior to Pasta, with the latter recovering significantly more senescence-inducing compounds (12.7% vs 18.9%; *p* = 0.025; Fisher’s exact test; Extended Data Fig. 4). A similar trend was observed for the prediction of reprogramming-promoting compounds (6.5% vs 16.9%; *p* = 0.142; Fisher’s exact test; Extended Data Fig. 4). Taken together, this data showed that Pasta identified age-modulatory compounds more effectively than a conventional regression model.

Analysis of the mechanisms of action (MOAs) annotated for the CMAP compounds revealed 26 enriched MOAs among Aging compounds, including inhibition of topoisomerases, cyclin-dependent kinases (CDKs), mouse double minute (MDMs), and inosine 5′-monophosphate dehydrogenase (IMPDHs), consistent with DNA damage and growth arrest pathways (Fig. 4c; Supplementary Table 8). Six MOAs were enriched among Rejuvenating compounds (Fig. 4d), dominated by HDAC inhibition, in line with their known role in accelerating reprogramming^63,64^. Additional Rejuvenating MOAs included MAPK/ERK kinase (MEK) and glycogen synthase kinase 3 (GSK3) inhibition, common components of reprogramming cocktails^62^, as well as cardiac glycosides and proteasome inhibitors, both previously described as senolytics^68^. Such senolytics may rejuvenate transcriptomic profiles indirectly, through selective elimination of aged cells (see Discussion).

Analyzing compound effects across cell lines revealed both conserved and context-dependent responses. Proteasome inhibitors MG-132 and bortezomib consistently reduced age scores, while mitoxantrone increased them (Fig. 4e; Supplementary Table 9). In contrast, several HDAC inhibitors (e.g., trichostatin-A, vorinostat, and panobinostat) showed divergent effects, reducing age scores in most cell lines but increasing them in neuronal lines (NEU, NPC, SH-SY5Y), consistent with prior reports of trichostatin-A’s cytotoxicity in SH-SY5Y cells^69^. These results underscore the dual and context-dependent roles of HDAC inhibitors in promoting both reprogramming and senescence^63–67^.

Chemotherapeutics have been described as having pro-aging properties^56^. Interestingly, we observed that most approved anti-cancer drugs were enriched in our set of Aging compounds (Supplementary Note 1, Supplementary Tables 10-12, Extended Data Fig. 5). Additionally, we identified compounds that had a stronger pro-aging effect in cancer cell lines than in normal cell lines. Such candidates might have translational applications for more effective and lower toxicity anti-cancer therapies. Furthermore, we could show that increase in cellular age is most often decoupled from increase in cell death (Supplementary Note 2, Supplementary Tables 13-15, Extended Data Fig. 6 and 7). This indicates that applying aging clocks to perturbation datasets could lead to novel anti-cancer therapies that have evaded conventional cell viability screens.

Collectively, these results demonstrated Pasta’s ability to detect both aging- and rejuvenation-inducing compounds in the CMAP L1000 data, underscoring its cross-platform robustness and ability to reveal conserved and context-dependent age-regulatory mechanisms.

### Experimental validation of compounds with age-modulatory effects

To experimentally test Pasta’s predictive power in identifying cell-type-specific age-modulatory compounds, we selected two top-ranked hits, one predicted to increase and one to decrease cellular age. For each compound, two cancer cell lines were chosen: a predicted *responder* expected to show the age-modulatory phenotype and a *non-responder* predicted to remain largely unaffected (Supplementary Table 16).

We first examined pralatrexate, the fourth-ranked Aging compound among 30,920 in the CMAP dataset (Supplementary Table 7). In the responder A375 melanoma cells, pralatrexate induced clear signs of senescence, including cell enlargement and positivity for the Senescence-Associated β-Galactosidase assay (Fig. 5a). It increased CDKN1A (p21) and IL6 (SASP component) expression (Fig. 5b), reduced proliferation as indicated by lower EdU incorporation (Fig. 5c), and downregulated LMNB1 while upregulating p21 protein (Fig. 5d), all consistent with a *bona fide* senescence response. In contrast, the non-responder MDA-MB-231 breast cancer cells displayed only mild stress responses lacking key hallmarks of cellular senescence: minimal morphological changes yet detectable SA-β-Gal positivity (Fig. 5e), negligible changes in CDKN1A and IL6 mRNA levels (Fig. 5f), increased proliferation and EdU incorporation (Fig. 5g), and modest LMNB1 variation accompanied by upregulation of p21 protein levels (Fig. 5h).

**Figure 5.**
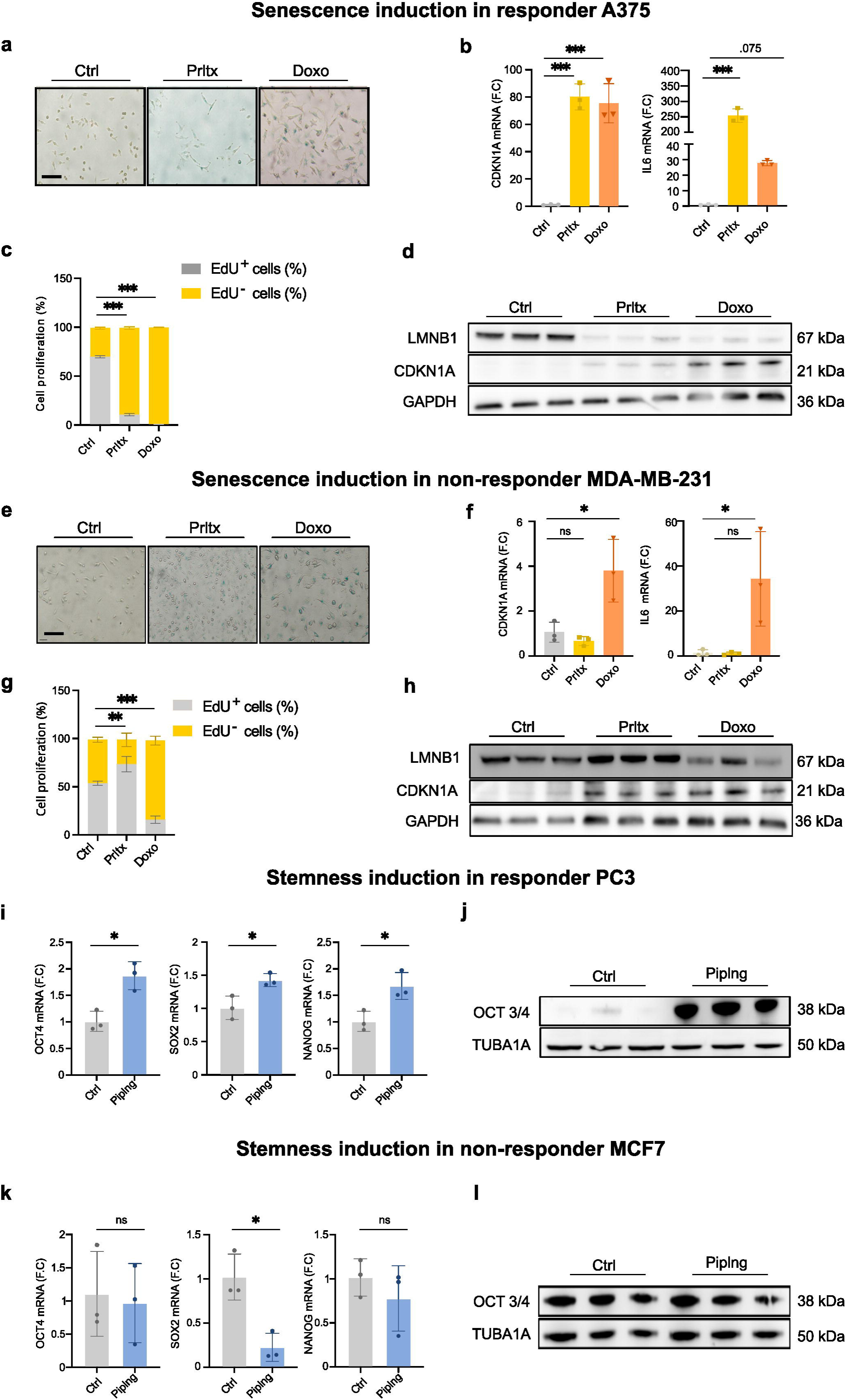
Experimental validation of pralatrexate as a senescence-inducer and piperlongumine as a rejuvenating agent. (**a-h**) Testing senescence-induction by pralatrexate in responder A375 and non-responder MDA-MB-231 cells. (**a,e**) Representative images from Senescence Associated Beta Galactosidase assays in A375 (**a**) and MDA-MB-231 (**e**) cells exposed to pralatrexate. Doxorubicin served as the positive control for senescence induction. Scale bar 10 micrometers. (**b,f**) qPCR analysis of CDKN1A and IL6 mRNA levels in A375 (**b**) and MDA-MB-231 (**f**) cells treated with pralatrexate or doxorubicin. N=3 biological replicates; *p< 0.05; *** p< 0.001; one-way ANOVA. (**c,g**) Cell proliferation by EdU incorporation in A375 (**c**) and MDA-MB-231 (**g**) cells after treatment with pralatrexate or doxorubicin. One representative experiment with three replicates per condition. ** p< 0.01; *** p< 0.001 (one-way ANOVA, calculated on EdU^+^ cells). (**d,h**) Immunoblotting showing LMNB1, CDKN1A and GAPDH protein levels in A375 (**d**) and MDA-MB-231 (**h**) cells after treatment with pralatrexate or doxorubicin. N=3 biological replicates. (**i-l**) Testing stemness-induction of piperlongumine in responder PC3 and non-responder MCF7 cells. (**i,k**) qPCR analysis of OCT4, SOX2 and NANOG levels in PC3 (**i**) and MCF7 (**k**) cells exposed to piperlongumine. * p< 0.05; Student’s t test. (**j,l**) Immunoblotting showing OCT3/4 and alpha-tubulin protein levels in PC3 (**j**) and MCF7 (**l**) cells treated with piperlongumine. Abbreviations: Prltx, pralatrexate; Doxo, doxorubicin; Piplng, piperlongumine; ns, non-significant.

Next, we evaluated Piperlongumine, ranked twentieth among 30,920 rejuvenating compounds (Supplementary Table 7). In the responder PC3 prostate cancer cells, piperlongumine enhanced expression of the pluripotency factors OCT4, SOX2, and NANOG and increased Oct4 protein abundance (Fig. 5i,j), indicating a transcriptional shift towards a youthful, stem-like state. In contrast, non-responder MCF7 breast cancer cells showed none of these phenotypes (Fig. 5k,l).

Together, these findings confirm that Pasta accurately predicts cell-type-specific age-modulatory outcomes, validating pralatrexate as a senescence-inducing and piperlongumine as a stemness-promoting compound.

### Systematic identification of age-increasing and rejuvenating genetic perturbations in the CMAP L1000 dataset

To uncover genetic drivers of cellular aging, we analyzed gene knockdowns, knockouts, and overexpression profiles in the CMAP L1000 dataset. Across cell lines, 841 gene perturbations (GPs) were significantly associated with higher age scores (*Aging GPs*) and 54 with lower age scores (*Rejuvenating GPs*).

The top two Aging GPs were overexpression of KRAS and BRAF (Fig. 6a; Supplementary Table 17). Furthermore, overexpression of three additional genes in the mitogen activated protein kinase (MAPK) pathway (RIT1, SOS1, and MAPK1) were in the top 12 pro-aging GPs. This result is consistent with the tumor-suppressive mechanism of oncogene-induced senescence (OIS), which is commonly triggered by hyperactivation of the MAPK pathway^70^. The third best Aging GP was knockout of CCNA2, consistent with a prior study reporting potent senescence induction by CCNA2 depletion^71^. The fourth and fifth best Aging GPs were knockout of the cyclin CCNA2 and knockdowns of ERG and MYC. Interestingly, even though ERG and MYC are also oncogenes, their overexpression does not seem to trigger OIS but instead repress it^72,73^. Accordingly, MYC depletion has been shown to promote senescence^73,74^.

**Figure 6.**
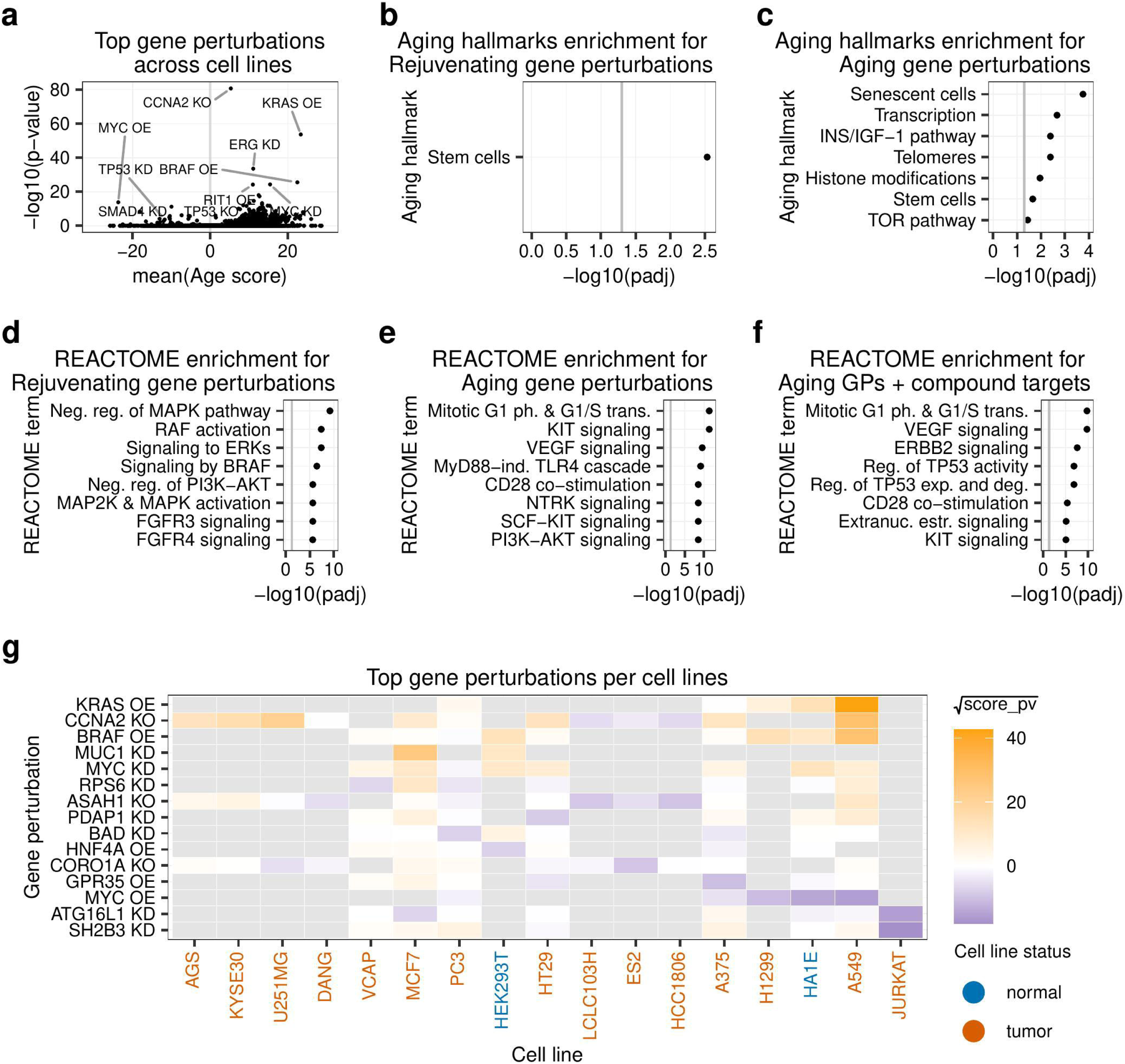
Pasta identifies age-modulatory gene perturbations. (**a**) Volcano plot displaying the most significant Rejuvenating gene perturbations (GPs) with negative age scores and Aging GPs with positive age scores. (**b-f**) Enrichment analysis using the Hallmarks of Aging^79^ (**b-c**) and REACTOME (**d-f**) gene sets for Rejuvenating GPs (**b, d**), Aging GPs (**c, e**), and Aging GPs that are also compound targets of significant Aging compounds (**f**). Significant results from the three GP types (OE, KD, and KO) were pooled, with overexpression results inverted (i.e., MYC OE was included in the Aging GP set). Grey lines mark the adjusted p-value cutoff of 0.05. (**g**) Heatmap of GPs by cell lines showing the most significant age scores (see Methods). Colors indicate the square root of a metric balancing significance and effect size: *-log_10_(adjusted p-value) × mean(age score)*. Missing values are shown in grey.

The top Rejuvenating GP was again connected to MYC – namely its overexpression – consistent with its key role in promoting stemness^75^ and repressing OIS^73^ (Fig. 6a; Supplementary Table 17). The second best Rejuvenating GP was SH2B3 knockdown, which aligns with the reported improvement in hematopoietic stem cell function upon Sh2b3 knockdown in *Fancd2^−/−^* mice^76^. The fourth best Rejuvenating GP was SMAD4 knockdown, which is consistent with SMAD4’s role in repressing self-renewal of neuroblastoma stem cells^77^. Finally, TP53 knockout and knockdown ranked as the 12^th^ and 18^th^ best Rejuvenating GPs. This result is consistent with the central tumor-suppressive role of TP53 and its mutation in 50-60% of human cancers^78^.

We then systematically characterized the significant age-modulatory GPs beyond the top hits. To this end, we inverted overexpression effects to pool all GP results and conducted gene set enrichment analysis using two complementary databases. The Aging Hallmark gene sets from the Open Genes database^79^ were used to explore aging signatures, and the REACTOME database^80^ gene sets were used to comprehensively study detailed cellular pathways (Fig. 6b–f; Extended Data Fig. 8a-f; Supplementary Tables 18 and 19). Among Rejuvenating GPs, the only enriched Aging Hallmark^79^ term was “stem cells” (Fig. 6b), while among Aging GPs the most enriched term was “senescent cells” (Fig. 6c). Aging GPs were also enriched for transcriptional regulation, insulin/IGF-1, mTOR, and telomeres pathways, all of which are related to aging and its regulation. In REACTOME, Rejuvenating GPs were enriched in RAF/MEK/ERK and FGFR signaling (Fig. 6d), whereas Aging GPs were enriched in pathways linked to cell cycle progression, receptor tyrosine kinase activation (KIT, VEGF, NTRK), PI3K/AKT signaling, and immune activation (TLR4, CD28) (Fig. 6e). To get a consolidated list of age-modulatory perturbations, we identified genes for which both genetic perturbations and compounds targeting them produced significant age effects in the same direction (Extended Data Fig. 8g). These genes were enriched in p53-dependent cell-cycle control and growth-factor signaling (VEGF, ERBB2, KIT, extranuclear estrogen) (Fig. 6f).

Next, we identified GPs with strong cell-line-specific age effects. MYC overexpression repeatedly produced rejuvenation, while MYC knockdown, CCNA2 knockout, and BRAF/KRAS overexpression increased cellular age (Fig. 6g; Supplementary Table 20). Others, such as SH2B3 knockdown and ASAH1 knockout, could either increase or decrease cellular age depending on context, underscoring the cell-type specificity of age regulation.

In summary, these findings map the genetic architecture of transcriptomic age modulation, revealing broad regulators like BRAF and MYC as well as context-dependent determinants of cellular age.

### Defining the molecular determinants of the propensity of cells to respond to age-modulatory perturbations

While we previously revealed the pathways that drive transcriptomic age prediction (Fig. 2) and that induce age-modulation (Fig. 4), it remains unclear which pathways determine the heterogeneity in responses of various cell lines to the same pro-aging or rejuvenating cues. We addressed this question by leveraging the extensive CMAP L1000 and DepMap datasets, by identifying molecular features that govern a cell’s propensity to age or rejuvenate. *Aging and rejuvenation propensities* were computed for 19 cell lines by averaging cell line-specific age scores of globally significant age-modulatory perturbations (see Methods). This analysis revealed marked heterogeneity (Fig. 7a), with MDA-MB-231 showing the highest rejuvenation propensity (−14.2 vs. −9.6 in MCF10A), consistent with its role as a model for aggressive, triple-negative breast cancer^81^.

**Figure 7.**
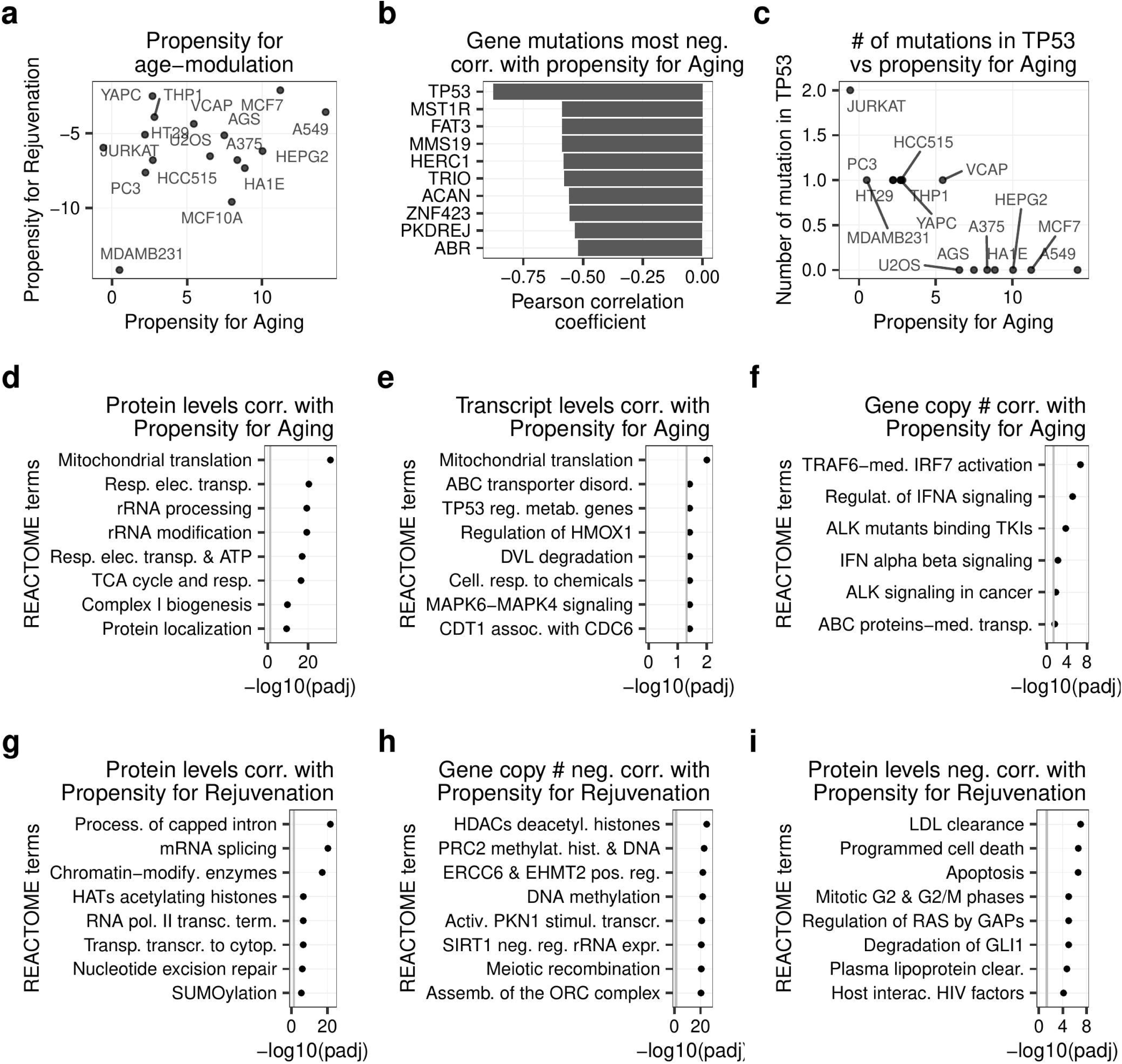
Pasta uncovers molecular determinants of cellular propensity for aging and rejuvenation. (**a**) Propensity of various cell lines to shift toward older or younger states following exposure to age-regulatory perturbations (see Methods). (**b**) Gene mutation counts most strongly negatively correlated with the propensity for Aging. (**c**) TP53 mutation count versus propensity for Aging. (**d-i**) REACTOME pathway enrichment for genes whose protein abundance (**d, g, i**), mRNA expression (**e**), and copy numbers (**f, h**) exhibit an absolute PCC higher than 0.4 with the propensity of cells for Rejuvenation (**d, e, f**) and Aging (**g, h, i**). The grey lines mark the adjusted p-value cutoff of 0.05.

When aging propensity was correlated with the number of non-silent mutations, TP53 emerged as the only significantly associated gene (PCC = 0.875; Fig. 7b; Supplementary Table 21). Cell lines harboring two TP53 mutations (i.e., JURKAT) had negative aging-propensity scores, those with one mutation scored below six, and TP53 wild-type lines scored higher (Fig. 7c). These results align with TP53’s established role as a tumor suppressor that enforces senescence in highly proliferative cells^82,83^ and suggest novel potential tumor suppressor genes (Fig. 7b). Consistent with this idea, top significant genes in Fig. 7b included the known tumor suppressor genes FAT3^84^, HERC1^85^, and ABR^86,87^.

To identify pathways influencing aging and rejuvenation propensities, we performed REACTOME enrichment analysis on proteins, transcripts, and gene copy numbers correlated with these traits (|PCC| > 0.4, Fig. 7d–i; Extended Data Fig. 9; Supplementary Table 22). Proteins and transcripts associated with aging propensity were both enriched in mitochondrial translation (Fig. 7d,e), underscoring mitochondrial activity as a central determinant of age-increase. This echoes recent findings linking p53, mitochondrial translation, and the SASP (see Discussion)^88^. Copy number–based enrichments included interferon-related pathways (Fig. 8f), consistent with type I interferons’ role in SASP^89^, DNA damage–induced^90^ and oncogene-induced senescence^91^, as well as their pro-aging effects^92^.

Rejuvenation propensity correlated with proteins involved in mRNA splicing, histone acetylation, and chromatin modification (Fig. 7g). At the copy number level, HDAC and PRC2 pathways showed the strongest negative associations (Fig. 7h). These findings align with evidence that HDAC inhibition promotes reprogramming^63,64^ and that PRC2 supports pluripotency^93^ and functions as a robust biomarker of aging and rejuvenation^94–96^. Proteins negatively correlated with rejuvenation propensity were enriched in apoptosis and checkpoint modules (Fig. 7i). This result is consistent with the finding that elevated expression of the Cdkn2a/Arf tumor-suppressor locus, which regulates senescence and apoptosis, constitutes a barrier for cellular reprogramming^97^.

Together, these results delineate the molecular architecture of cellular age control, where mitochondrial translation and chromatin regulation act as opposing axes.

## Discussion

In this study, we introduce Pasta, a versatile multi-tissue transcriptomic aging clock that robustly predicts relative age across diverse human and mouse transcriptomic datasets. Pasta outperformed existing transcriptomic clocks, captured cellular senescence and stemness states, and identified genetic and chemical modulators of aging. It is available as a user-friendly, open-source R package to support broad use in aging research.

A key strength of Pasta lies in its adaptability to diverse transcriptomic platforms, which was achieved by using an age-shift learning framework trained on heterogeneously processed RNA-Seq and microarray datasets. This design enhanced cross-dataset robustness and allows users to input gene-expression count data directly, without cross-dataset normalization or batch correction. Pasta achieved strong accuracy across tissues and data types and relied on biologically relevant features that produced more meaningful predictions than conventional regression models, positioning age-shift learning as an effective approach for predicting relative age.

Pasta’s gene coefficients and enrichment analyses converge on the DNA damage response and its downstream effector, p53 signaling, as key pathways linked to transcriptomic aging. Core predictors such as CDKN2A/p16 and ZMAT3, both canonical p53 targets, underscore this mechanistic grounding. Downstream perturbation analyses reinforced these links: topoisomerase inhibitors, which induce DNA damage, and CDK inhibitors, which block the cell cycle, were the two most enriched mechanisms of action among aging-associated compounds, and pathways related to G1/S transition and TP53 regulation were consistently enriched across chemical and genetic pro-aging perturbations. Notably, recent work has proposed the DNA damage response as a fundamental “counting unit” of aging clocks, in which DDR, cell-cycle progression, and senescence act as interconnected temporal layers of cellular timekeeping^98^. Within this framework, the transcriptomic aging patterns captured by Pasta may reflect features emerging from these distinct but coordinated biological timescales.

Pasta showed strong predictive power for identifying cellular senescence, stemness, and age-related dynamics such as reprogramming and differentiation, despite not being trained on senescence or stem cell data. This suggests that transcriptomic aging clocks capture local cellular aging effects along a continuum from stemness to senescence. Future studies could assess whether aging clocks trained on other molecular layers, such as DNA methylation or proteomics, display similar behavior.

Pasta’s age scores correlated significantly with tumor grade and survival across multiple cancer types, supporting its potential use for patient stratification. Moreover, Pasta’s age scores showed a stronger association with tumor grade than donor age, suggesting that the model captures biological aging signatures beyond chronological age.

Analysis of the CMAP data showed marked enrichment of senescence-inducing perturbations in the Aging group and of reprogramming- and stemness-associated perturbations in the Rejuvenating group. Notably, Aging perturbations (259 compounds, 841 GPs) greatly outnumbered Rejuvenating ones (59 compounds, 54 GPs), consistent with a recent report^23^, suggesting that diverse perturbations can accelerate aging, whereas rejuvenation requires higher precision. Perturbation effects were largely consistent across both cancer and normal (as defined in the CMAP cell annotation file) cell lines (Fig. 4e, 6g, Supplementary Tables 9, 16), indicating conservation of aging and rejuvenation pathways in health and disease. Interestingly, chemotherapeutics were strongly enriched in pro-aging compounds. These results are consistent with the view that many cancer drugs act by inducing senescence^99^ and increasing biological age of both normal and cancer cells^16^.

Despite a lot of consistency with the literature, some of our results were less expected. For instance, proteasome inhibitors such as MG-132 and bortezomib have been shown to transiently enhance pluripotency markers early after treatment^100^ but to ultimately suppress them at later stages^101^. Given that we identified them as rejuvenating compounds argues that Pasta might particularly capture the earlier effects of perturbations, consistent with CMAP’s short exposure duration (∼24 hours). Furthermore, these proteasome inhibitors as well as several other of our top rejuvenating hits, including piperlongumine and the structurally related CT-200783, as well as cardiac glycosides such as ouabain—consistent with recent reports^23,102^— and HDAC inhibitors, have previously been described as senolytic agents^68^. Here one may speculate that these compounds do not intrinsically rejuvenate cells but rather eliminate aged or damaged cells from the population.

Among the chemical perturbations identified, pralatrexate emerged as a previously unrecognized senescence-inducing compound. This dihydrofolate inhibitor, approved by the FDA in 2009 for relapsed or refractory peripheral T-cell lymphoma^103^, ranked fourth for senescence induction across cell lines and was validated experimentally. Its senescence-inducing effect mirrors those of other dihydrofolate inhibitors such as methotrexate^104–106^ (ranked 98th), pemetrexed^107^, and pyrimethamine^108^. Notably, pralatrexate ranked first for preferentially inducing senescence in cancer cells, suggesting potential therapeutic selectivity. These results highlight the ability of transcriptomic aging clocks such as Pasta to reveal latent pharmacological properties of established drugs.

We observed heterogeneity in cell line responses to age-modulatory perturbations, such as HDAC inhibitors having pro-aging or pro-rejuvenating effects depending on the cell lines. To better understand the origins of this heterogeneity, we integrated CMAP and DepMap data to identify molecular determinants of the propensity of cells to respond to age-modulatory cues. Transcriptomic and proteomic analyses showed that pro-senescence responses depend strongly on mitochondrial translation, consistent with links between mitochondrial metabolism, reactive oxygen species, and senescence^109^. Mitochondrial translation promotes biogenesis and reactive oxygen species production^110^, driving p21-mediated senescence^111^, while p53 enhances this process to sustain cytokine output through the SASP^88^. Increased type I interferon gene copy numbers also correlated with stronger aging responses, in line with the pro-aging role of interferon signaling^92,112^. These findings suggest that mitochondrial translation and interferon pathways sensitize cells to senescence via oxidative and inflammatory mechanisms.

Rejuvenation was instead shaped by mRNA splicing and epigenetic factors. Reduced PRC2 and HDAC gene copy numbers heightened sensitivity to rejuvenating stimuli, consistent with PRC2’s role as a universal biomarker of aging and rejuvenation^94^ and with HDAC inhibition facilitating reprogramming^113^. Together, these results delineate key molecular pathways underlying propensity for cellular aging and rejuvenation and highlight potential targets for therapeutic modulation.

To conclude, Pasta is a biologically grounded and versatile transcriptomic aging clock with broad utility, spanning fundamental research on human and mouse aging, senescence, and stem cells to clinical diagnostics and therapeutic discovery. By enabling systematic screening of perturbation datasets, it extends the use of conventional aging clocks toward data-driven inference of determinants of cellular biological age. Given the rapid expansion of perturbation datasets, this framework provides a scalable platform for translational research in cancer^16^, neurodegeneration^17^, regeneration^18–20^, and age-related interventions (Extended Data Fig. 10).

## Methods

### Construction of the aging transcriptomic database

GTEx Protected Access Data version 8 (17,382 samples) was used to obtain precise age information^37^. We discarded 1,674 samples with a RIN score below 6. An additional 1,137 samples originating from gender-specific tissues (cervix, fallopian, uterus, prostate, ovary, testis, vagina), tissues with low sample sizes (e.g., bladder, 12 samples), or non-relevant sources (e.g., *cells_ebv_transformed_lymphocytes*) were further discarded. Raw or normalized count matrices were retrieved in R using the package GEOquery^114^ for GEO^22^ datasets and using the package ExpressionAtlas^38^ for Expression Atlas^115^ datasets. The biomaRt R package^116^ was used to convert gene or probe IDs into Ensembl gene IDs. Custom scripts were used to parse health status, age, and tissue metadata. Overall, 21 datasets with data from healthy donors were identified. These datasets contained 17,212 samples in total, including 14,571 from GTEx, and 8,113 genes in common, which were also included in the landmark or best inferred gene sets in the CMAP L1000 dataset^13^ and were used for further analyses. A rank transformation was applied to each sample using base R’s *rank* function with the parameter *ties.method = ‘average’*.

### Generating sample pairs

For age-shift models, pairs of samples from the same dataset were generated randomly and then filtered. Sample IDs were shuffled 200 times for the training set and 2000 times for the testing set within each dataset to generate IDs for both the first and the second samples in the pairs. Then, pairs with an age difference below the cutoff, involving non-healthy individuals, or representing duplicate pairs (i.e., sample 1 and sample 2 swapped) were excluded. Datasets with more than 1.5 times as many pairs as samples were randomly downsampled to maintain a ratio of 1.5 pairs per sample. Then, the ranked gene expression values were subtracted between sample 1 and sample 2 and used as input to train the models. The output variable was binary, indicating whether sample 1 was younger or older than sample 2.

### Training and evaluating models

Ridge-regularized generalized linear models were built using 10-fold cross-validation to identify the optimal lambda parameters. This was done using the function *cv.glmnet* from the glmnet package^117^, and using the following parameters: *s_lambda = ‘lambda.min’*, and *type.measure = ‘mse’* for regression models or *type.measure = ‘deviance*’ for classification models. Models were compared using a leave-one-dataset-out (LODO) cross-validation strategy. The age effects of the classification models (traditional and age-shift classifiers) refer to the log-odds obtained using the *predict* function from the glmnet package with parameter *type = ‘link’*. The Pearson correlation coefficients of the predicted ages (for the regression model) or age effects (for the classification models) with the true ages in the left-out datasets were used as the metric to compare models. To convert the classifiers’ predictions into age differences, referred to as age scores, they were multiplied by a scaling factor specific to each classifier. These scaling factors were obtained by applying the cross-validation models to a set of pairs with balanced age differences in the left-out datasets. Specifically, 1000 pairs were generated per dataset which were then filtered to retain 20 pairs in each of 13 bins representing 10 year age differences. Age predictions were made and results across all datasets were aggregated. Finally, the scaling factor was defined as the beta coefficient from a linear regression of age_difference on age_effect.

### Comparing Pasta with MultiTIMER

The MultiTIMER model cannot be used directly, as it requires query samples to be normalized alongside the training samples^30^. As a result, it only functions with samples available on ARCHS4^9^. We re-ran the MultiTIMER training code using ARCHS4 version 2.1, incorporating 14 test datasets —8 from the LODO analysis and 6 novel datasets—alongside the MultiTIMER’s training samples to ensure joint batch correction. LODO datasets were evaluated using Pasta’s LODO performance, while the novel datasets were assessed using predictions from the final Pasta model.

### Extending Pasta for application to mouse samples

Orthologous gene pairs between human and mouse were obtained from the Human Gene Database HGD^118^. The table was filtered to retain only one-to-one orthologues. For each mouse dataset, gene symbols or probes were converted to Ensembl gene identifies. Genes were then restricted to those with one-to-one human orthologues and renamed using human gene names. This yielded 3292 genes, of which 1,600 overlapped with Pasta’s predictive genes. Missing genes required for the Pasta age model were imputed with the dataset median, and rank-transformations were conducted before applying the age model.

### Conducting gene set enrichment analysis

GSEA was performed using the GSEA function from the *clusterProfiler* R package^119^. Gene sets were retrieved via the msigdbr function from the *msigdbr* package^120^, aggregating sets from the CGP, KEGG, REACTOME, and MIRDB subcategories. Model coefficients, sorted by decreasing magnitude, were used as input. The *TERM2GENE* argument was set to the aggregated gene sets, with all other parameters left at default.

### Analyzing senescence and stem cell datasets

Supplementary Table 1 from the SenCID manuscript^121^ was used to identify samples from senescent, proliferative, and quiescent cells. Custom scripts were used to identify GSM IDs for each sample. This resulted in 308 of 603 samples which could be downloaded and analyzed. For stem cell datasets, samples were manually annotated. Data were downloaded mainly from GEO using Pasta’s functions, *getting_geo_count_mat_for_age_prediction* and *getting_GEO_ES_for_age_model*, as well as with custom scripts. The Pasta R package was used to make age predictions. AUC-ROC values were computed using the *auc* function from the pROC R package^122^.

### Analyzing TCGA data

Histological tumor grades were obtained from Ping et al^123^ for samples from BRCA patients and from Liu et al^124^ for samples from patients with other cancer types, as described in Zheng et al^57^. Only cancer types with histological tumor grades ranging from G1 to G4 were retained, excluding BLCA samples. The TCGAbiolinks R package was used to download gene expression data and obtain age and overall survival information^125^. Only tumor samples were used for all analyses. The Pearson product-moment correlation coefficient test, implemented in base R’s cor.test function, was used to compute p-values and confidence intervals for the correlation between Pasta scores or age in days with tumor grades (from 1 to 4). P-values were adjusted for multiple testing using the False Discovery Rate method implemented via base R’s *p.adjust* function with *method = ‘fdr’*. Survival analyses were conducted by fitting Cox proportional hazards regression models using the *coxph* function from the R survival package^126^. The predictor variable was either standardized Pasta age scores or standardized age in days. Hazard ratios and their confidence intervals were obtained by exponentiating the model’s coefficients and their confidence intervals. Kaplan-Meier survival analyses were performed using the *survfit* function from the R survival package, with Pasta’s tertiles as the predictor variable.

### Analyzing CMAP L1000 data – Preprocessing data and predicting age scores

Raw level 3 data and metadata files were obtained from the https://clue.io website, and MD5 checksums were verified. Several CMAP plates had incorrect or missing annotations, which were corrected using custom scripts. Plates with missing treatment information or fewer than three control samples were excluded from all analyses. The perturbation metadata table was reduced from 3,027,596 to 3,017,946 entries after filtering. Missing mechanisms of action (MOAs) were manually annotated for 60 compounds. Missing compound names were manually annotated for seven compounds that had only BRD (Broad) IDs available. All data were split into individual *.rds* files by plate to facilitate parallel computation with reduced memory usage. For each sample, its final age score was defined as its raw age score minus the mean age score of control samples on the same plate. Control samples were defined as DMSO-treated for compound perturbations and untreated for genetic perturbations.

### Analyzing CMAP L1000 data – Annotating senescence- and reprogramming-related compounds

Senescence-inducing compounds were obtained from Table 1 in Petrova et al^61^. Reprogramming-promoting compounds were obtained from Supplementary Table 1 in Knyazer et al^62^. Due to the large number of compounds listed, only those used in at least 3 studies were selected. The selected compounds were manually checked for their presence in the CMAP dataset. In total, 31 senescence-inducing compounds and 11 reprogramming-promoting compounds were identified. The MOA in CMAP for each of these compounds was identified. MOAs for compounds present in both sets were annotated as ‘both’. This resulted in 5 MOAs for the ‘reprogramming’ group, 11 MOAs for the ‘senescence’ group, and 2 MOAs for the ‘both’ group. All CMAP compounds belonging to these MOAs were annotated accordingly.

### Analyzing CMAP L1000 data – Statistical analysis

Two-sided t-tests were used to assess whether age scores significantly differed from zero for each perturbation, either across all cell lines or within each individual cell line. All dosages, incubation times, and BRD IDs for a given perturbation were pooled to increase statistical power. Bonferroni correction was applied to account for multiple testing. For GPs, a two-stage approach was employed to mitigate noise associated with variability in genetic construct quality. In the first stage, significance was tested for each BRD ID. In the second stage, significance was tested for each GP using only its top two BRD IDs with the highest absolute average score_pv values. Two metrics were created to combine statistical significance with effect size: *score_pv = −log10(p-value) * mean(age score)*, and *score_pa = −log10(adjusted p-value) * mean(age score)*. Significant results were defined as entries with a score_pa greater than or equal to −log10(0.05). Over-representation of MOAs among the significant entries, relative to other entries, was assessed using Fisher’s exact test, implemented in base R’s fisher.test function with *alternative = ‘greater’*. FDR correction was applied to account for multiple testing. Analysis of cancer-specific pro-aging compounds was conducted using one-sided two-sample t-tests to assess whether the mean age in tumor cell lines was greater than that in normal (as defined in the CMAP cell annotation file) cell lines. Only compounds with at least two normal and two tumor cell lines and a mean tumor age effect greater than two were selected. The score_pa and score_pv metrics were derived as described above but using the difference in mean age effect between tumor and normal cell lines as the measure of effect size.

### Analyzing CMAP L1000 data – Selecting perturbations and cell lines to display in the heatmaps

Perturbation and cell line selection was performed separately for the Aging and Rejuvenating groups. The perturbation-by-cell line results tables were sorted by decreasing score_pa for the Rejuvenating group and by increasing score_pa for the Aging group. The top 40 perturbations were selected for each group, with a maximum of one perturbation per cell line, which ensured that we displayed the most significant results from our study. For each selected perturbation, the top 5 cell lines were selected.

### Analyzing CMAP L1000 data – Pathway enrichment analysis

Significance results from overexpressed genes were inverted to generate a pooled set of GPs with consistent directional effects per gene. The Aging Hallmarks gene sets were obtained from Open Gene^79^. Terms were standardized to the noun forms as these were used for creating the original gene sets. REACTOME gene sets were obtained from the Human MSigDB Collections^46^, using the msigdbr R package^120^. Only REACTOME pathways with 300 genes or fewer were retained (1,565 of 1,615). Over-representation analysis was performed using the *enricher* function from the clusterProfiler R package^119^ with the parameter *universe* set to all the unique genes in the input database.

### Analyzing CMAP L1000 data – Analysis of anticancer drugs

Anticancer drugs were obtained from the CancerDrugs_DB^127^ (build date: 28/11/24). We selected drugs approved by the FDA, the EMA, or European national agencies, which retained 311 of 321 drugs. We then kept 132 drugs that were present in the CMAP L1000 dataset. For each cancer, over-representation of its anticancer drugs among the compounds from a given age set (Aging, Rejuvenating or Other), relative to other compounds, was assessed using Fisher’s exact test, implemented in base R’s fisher.test function with *alternative = ‘greater’*. FDR correction was applied to account for multiple testing.

### Analyzing CMAP L1000 data – Identifying consistent compound target / gene perturbation results

To identify significant GPs whose age effects are consistent with the targets of significant compounds, we computed the mean score_pa for each gene, using the pooled GP significance table, and for each compound target. Compounds without annotated targets were excluded from this analysis. For compounds with multiple annotated targets, each target was analyzed separately. This resulted in mean score_pa values for 7,638 genes and for 868 targets of compounds. The intersection of these two sets yielded 813 entries, of which 591 had a consistent directionality of effect and were retained. 47 of these had non-zero score_pa values in both sets and were retained. Finally, only perturbation pairs with a min score_pa below 0.05 were retained to include only genes significant in both genetic and chemical perturbations. This resulted in 1 gene for the rejuvenation set (MAP2K1) and 14 genes for the Aging set.

### Analyzing DepMap data – Data sources

DepMap data were obtained using the depmap R package^128^. Cell line metadata was retrieved using the *depmap_metadata* function. Drug, CRISPR and RNAi sensitivity data were obtained using the *drug_sensitivity_21Q2*, *depmap_crispr*, and *depmap_rnai* functions, respectively. Copy number, gene expression, protein abundance, and mutation data were obtained using the *depmap_copyNumber*, *depmap_TPM*, *depmap_proteomic*, and *depmap_mutationCalls* functions, respectively.

### Analyzing DepMap data – Dependency scores

The significance of age scores for compounds was computed for each BRD ID and cell line, using the same approach as described above (see Statistical analysis). CMAP age score significance results and DepMap dependency scores were then merged by BRD IDs for compounds (41,199 overlapping entries), and by gene names for CRISPR (46,752 overlapping entries) and RNAi (27,401 overlapping entries). The associations between dependency scores and perturbations (compound, CRISPR or RNAi) were assessed by fitting linear models with dependency score as the dependent variable and mean age score as the independent variable. The association between dependency scores and gene sets (MOAs for compounds and REACTOME pathways for CRISPR) was tested by fitting linear models with dependency score as the dependent variable and both mean age score and perturbation name as the independent variables. P-values were adjusted for multiple testing using the Bonferroni correction. This analysis resulted in the identification of 2 MOAs, 2 compounds, 9 REACTOME pathways and no gene perturbations whose mean age scores were significantly associated with dependency scores at an adjusted p-value significance cutoff of 0.05. The relatively low number of replicates per gene perturbation (median 9, 3^rd^ quartile 12) could explain the lack of significant entries for this analysis.

### Analyzing DepMap data – Computing cell lines’ propensity for Aging and Rejuvenation

The compound-by-cell line and GP-by-cell line age score significance tables were merged and filtered to retain only perturbations with significant age scores across all cell lines. To derive robust conclusions, the table was further filtered to retain only entries from 19 cell lines in which at least 25 significant perturbations had been tested for both the Rejuvenation and the Aging groups. Propensity scores were then computed as the mean age score of all significant perturbations in the Rejuvenation and Aging groups.

### Analyzing DepMap data – Correlating propensity scores with molecular features

The significance of the association between gene mutation and propensity scores was assessed using the Pearson product-moment correlation coefficient test, implemented in base R’s *cor.test* function. P-values were corrected for multiple testing using the Bonferroni method. Of 1,724 genes, only TP53 showed a significant association. PCCs between cell lines’ propensity for Aging or Rejuvenation and protein expression levels, log-transformed gene copy numbers, or gene expression (in transcripts per million) were computed for each molecule (e.g., gene or protein). Molecules with an absolute PCC higher than 0.4 were analyzed using ORA with the REACTOME database, as described above (see Pathway enrichment analysis).

### Cell Culture

A-375 (human melanoma) cells were kindly provided by Dr. Aishe Sarshad (University of Gothenburg, Sweden) and Dr. Alexander Espinosa (Karolinska Institute, Sweden). MDA-MB-231 (human breast cancer) cells were provided by Pr. Kirsty Spalding (Karolinska Institute, Sweden). PC3 cells were kindly provided by Dr. Yvonne Ceder (Lund University). Jurkat cells were purchased from ATCC. Cell lines were maintained in standard DMEM (Gibco). All media were supplemented with 10% heat-inactivated fetal bovine serum (FBS; Gibco) and 1% antibiotics (penicillin/streptomycin 100 U/mL; Gibco). Cells were maintained in a humidified incubator at 37°C and 5% CO2. Cells were tested monthly for *Mycoplasma* contamination using MycoAlert^TM^ Mycoplasma detection kit (Lonza^TM^ LT07-4118-11630271), and only negative cells were used for experiments. For the assessment of pro-aging or rejuvenating effects, cells were incubated in the presence of Pralatrexate (MedChemExpress, #HY-10446, 100 nM in DMSO) or Piperlongumine (MedChemExpress, #HY-N2329, 10 μM) for 72 hours. The DNA damage-inducing agent doxorubicin (D1515, Sigma-Aldrich, 100 nM) was used as a positive control for senescence induction.

### SA**β**G Assay

Cells were washed with PBS, fixed with 0.2% glutaraldehyde for 10 minutes, washed with PBS, and incubated overnight at 37°C with a staining solution containing 1 mg/mL X-Gal (Calbiochem, #203782-1GM) prepared in dimethylformamide (DMF; Sigma-Aldrich, D4551) at pH 6. Cells were then washed in PBS and visualized using an Olympus IX73 brightfield microscope.

### Cell Proliferation analysis

Cell proliferation was assessed by EdU incorporation by Click-iT™ Plus EDU Alexa Fluor™ 647 assay (C10634, Thermo Fisher Scientific) as described by the manufacturer. EdU positivity was assessed by Flow Cytometry using a BD FACSCanto II (BD) at the Biomedicum Flow Core Facility. FACS data were analyzed using the FlowJo software.

### Immunoblotting

For immunoblotting, proteins extracted by cellular lysis in RIPA buffer were run on 4–12% Bis-Tris acrylamide gels (#NP0322, Thermo Fisher Scientific) and electrotransferred to 0.2Cμm polyvinylidene fluoride (PVDF) (#11704156, Biorad) using Trans-Blot Turbo Transfer System (BioRad). Non-specific binding sites were saturated by incubating membranes for 1 h in 0.05% Tween 20 (#P9416, Sigma Aldrich) v-v in Tris-buffered saline (TBS) supplemented with 5% BSA (#10735078001, Sigma-Aldrich, w:v in TBS), followed by an overnight incubation with primary antibodies specific for Waf1/Cip1/CDKN1A p21 (#sc-6246, Santa Cruz Biotechnology), LaminB1 (12987-1-AP, Proteintech) and Oct3/4 (sc-5279, Santa Cruz Biotechnology). Equal protein loading was monitored with GAPDH (#2118, Cell Signaling) and alpha-tubulin (#2144, Cell Signaling) specific antibodies. Membranes were cut in order to allow simultaneous detection of different molecular weight proteins. Membranes were developed with suitable horseradish peroxidase conjugates followed by chemiluminescence-based detection with the Cytiva Amersham ECL Prime (#RPN2232, GE Healthcare) and the Amersham ImageQuant500 software-assisted imager (Cytiva).

### Gene Expression Analysis by RT-qPCR

Total RNA was extracted from cell samples using TRIzol reagent (Invitrogen, #15596018) according to the manufacturer’s instructions. Up to 1 μg of total RNA was reverse transcribed into cDNA using the iScript cDNA Synthesis Kit (Bio-Rad, #1725038). Quantitative real-time PCR was performed using GoTaq PCR Master Mix (Promega, #A6002) in a QuantStudio 5 Real-Time PCR System (ThermoFisher). The average expression of ACTB (for detection of senescence genes) or GAPDH (for detection of stemness genes) served as endogenous normalization controls. Primers used in this study are the following:

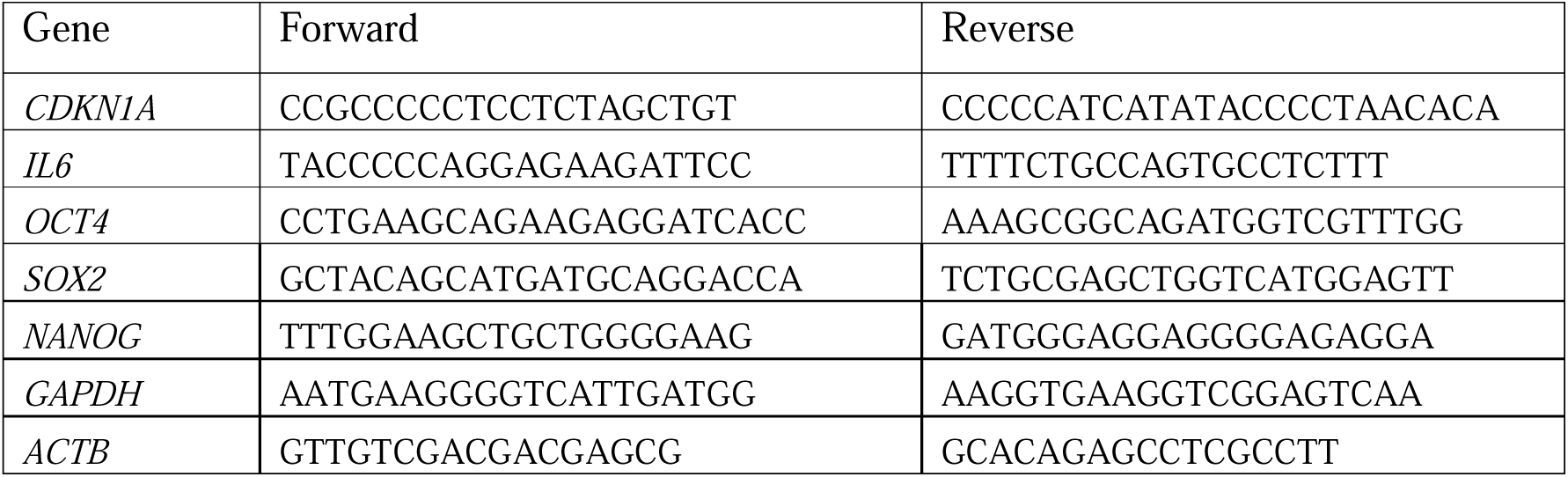

## Data availability

All datasets used to train and evaluate the Pasta model are publicly available from the GTEx Portal (https://gtexportal.org), the Gene Expression Omnibus (GEO; https://www.ncbi.nlm.nih.gov/geo/), and the Expression Atlas (https://www.ebi.ac.uk/gxa/home). Detailed accession numbers and metadata for each dataset are listed in Supplementary Supplementary Table 1. The Cancer Cell Line Encyclopedia (CCLE) and DepMap datasets were accessed via the depmap R package (https://depmap.org/portal/download/). The CMAP L1000 perturbation dataset was obtained from the Connectivity Map website (https://clue.io). The open source Pasta R package, including pre-trained models, functions for data download, processing, and age prediction, is publicly available on GitHub at https://github.com/jsalignon/pasta.

## Acknowledgements

We thank all members of the Riedel and Pietrocola laboratories for their valuable feedback. We acknowledge the GTEx Consortium, GEO, Expression Atlas, TCGA, CMAP, and DepMap teams for making their transcriptomic and functional genomics datasets publicly available, enabling this research. We thank Sara Hägg (Karolinska Institute) for providing early advice on validation of the transcriptomic aging clock. We thank Bader Zarrouki (AstraZeneca) for scientific discussions.

## Author contributions

J.S. performed all computational analyses, developed the R package, generated the computational figures and tables in the manuscript, and wrote the first draft of the manuscript. M.T., P.M.G., and E.R.D. conducted wet lab experiments. F.P., M.T., and P.M.G. generated wet lab figures and associated methods. J.S., H.A., F.P., and C.G.R. contributed to subsequent editing and revision of the manuscript. J.S., F.P., and C.G.R. conceived and supervised the study.

## Supplementary Figures legends

**Extended Data Fig. 1. Results of leave-one-dataset-out (LODO) analysis.**

(**a**) Comparative performance in predicting relative age, assessed with LODO, when considering only young (< 40 years old) and old (> 60 years old) individuals. (**b**) Detailed LODO results for the AS40/Pasta model when considering all individuals. The x-axis shows the AUC-ROC for predicting whether the first sample is younger than the second in each evaluation pair.

**Extended Data Fig. 2. Predicted versus true age differences for sample pairs using the AS40/Pasta LODO models.**

For each dataset, predictions were generated by a model trained on all other datasets. Each point corresponds to a sample pair with an age difference of at least 40 years.

**Extended Data Fig. 3. Predicted age scores versus true ages for all samples using the AS40/Pasta LODO models.**

For each dataset, predictions were generated by a model trained on all other datasets. Each point corresponds to one sample.

**Extended Data Fig. 4. Baseline regression model identifies fewer known age-modulatory compounds in L1000 data.**

Proportion of compounds annotated as senescence-inducing, pro-reprogramming, both, or neither for significantly age-increasing (Aging), age-decreasing (Rejuvenating) compounds, and other compounds (Other). Analyses were performed identically to Figure 4b but employed the baseline regression model from Figure 1 rather than the age-shift model.

**Extended Data Fig. 5. Chemotherapeutics broadly elevate transcriptomic age.**

(**a**) Cancers whose anticancer drugs are significantly enriched in age-modulatory compounds across all cell lines. Only cancers with at least 5 approved anticancer drugs according to CancerDrugs_DB^127^ were selected. 17 of 27 cancers were significant for the Aging group. The grey line marks the adjusted p-value cutoff of 0.05. (**b**) Top 10 compounds whose age scores are statistically higher in cancer cell lines than in normal cell lines (see Methods). (**c**) Mean age score in cancer and normal cell lines for the 10 compounds shown in (b).

**Extended Data Fig. 6. Relationship between compound-induced changes in cellular age and viability.**

(**a-c**) Yellow, blue, and black indicate compounds in the Aging, Rejuvenating, and Other groups, respectively. (**a**) Boxplots and violin plots of dependency scores for compounds by cell line in the Aging (175 entries), Rejuvenating (75 entries), and Other (40,906 entries) groups. (**b-c**) Mechanisms of action (**b**) and compounds (**c**) with the strongest associations between age scores and dependency. (**d**) Mean age score versus dependency for MDM inhibitors. Each point represents a compound in a given cell line.

**Extended Data Fig. 7. Relationship between gene perturbation-induced changes in cellular age and viability.**

(**a-b**) Yellow, blue, and black indicate gene perturbations (GPs) in the Aging, Rejuvenating, and Other groups, respectively. (**a**) Boxplots and violin plots of dependency scores for GPs by cell line in the Aging, Rejuvenating, and Other groups. Number of entries per set: Other (KD: 27,243; KO: 46,684 entries), Aging (KD: 152; KO: 55), Rejuvenating (KD: 6; KO: 13). (**b**) REACTOME pathways whose PCCs are most significantly associated with dependency scores in the knockout Aging group. No term was enriched in the Rejuvenating group. The grey lines mark the 0.05 adjusted p-value cutoff. (**c-d**) Mean age score versus dependency for GPs in the retinoic acid (RA) biosynthesis (**c**) and in the oncogene-induced senescence (**d**) pathways. Each point represents a gene knockout in a given cell line.

**Extended Data Fig. 8. REACTOME enrichment analysis of age-modulatory gene perturbations.**

Results for Rejuvenating knockdown (**a**), knockout (**b**), and overexpression (**c**), and for Aging knockdown (**d**), knockout (**e**), and overexpression (**f**). (**g**) Genes with non-zero adjusted p-value scores (score_pa) for their age-effect across cell types that showed consistency between the pooled gene perturbations and compound target analyses.

**Extended Data Fig. 9. REACTOME enrichment analysis of pathways significantly associated with cellular propensity for Aging and Rejuvenation.**

Results for genes whose copy number positively correlates with the propensity for Rejuvenation (**a**), proteins whose abundance negatively correlates with the propensity for Aging (**b**), and genes whose transcript levels positively correlate with the propensity for Rejuvenation (**c**) or negatively correlate with the propensity for Rejuvenation (**d**).

**Extended Data Fig. 10. Graphical summary of the study.**

